# Ports of Entry: Tracking Marine Invaders in Portuguese Marinas with DNA Metabarcoding from Coast to Archipelagos

**DOI:** 10.1101/2025.06.23.660658

**Authors:** Ana S. Lavrador, Inês Afonso, Paula Chainho, Ana Cristina Costa, João Paulo Medeiros, Manuela Isabel Parente, Paola Parretti, Patrício Ramalhosa, Ronaldo Sousa, Pedro E. Vieira, Filipe O. Costa, Sofia Duarte

## Abstract

Marine non-indigenous species (NIS), introduced primarily through shipping in marinas and ports, pose significant threats to coastal biodiversity and ecosystem functioning, requiring effective management strategies, especially for monitoring aimed at early detection. This study aimed to capture a comprehensive snapshot of the geographic variation of marine invertebrate assemblages and NIS occurrence, across 10 recreational marinas in Portugal (6 on the mainland - Viana do Castelo, Porto, Aveiro (2 marinas), and Lisbon (2 marinas) - and 2 in each archipelago, Madeira and the Azores). In each marina, hard substrates, zooplankton, and water samples (for eDNA) were collected in triplicate during summertime for metabarcoding profiling, adding up to a total of 111 samples. Using Illumina MiSeq sequencing of the mitochondrial cytochrome c oxidase subunit I (COI) gene (313 bp) and the V4 region of the small subunit 18S rRNA gene (18S, approximately 400 bp), we detected 645 species from 21 phyla, 40 of which were NIS (7 phyla). Only 5% of the species and 5% of NIS were detected in all marinas. The highest numbers of both exclusive native species and NIS were recorded in the Azores (100 and 7, respectively), and the lowest numbers of exclusive native species were recovered in Aveiro (39) while the lowest numbers of exclusive NIS were detected in the North and Aveiro regions (1 in each). A Principal Coordinate Analysis indicated a distinct separation among communities forming three main groups: 1) Viana do Castelo, Porto, and Aveiro; 2) Lisbon; and 3) Madeira and the Azores. Similarity between these groups is negatively correlated with geographic distance and overlaps with ecoregions previously identified within Lusitania. In the Azores 9 NIS were recorded for the first time, and one in Madeira. We also recorded a considerable number of potential range expansions of NIS, predominantly along the mainland coast, in both North (N) – South (S) or S-N directions, which shows promising results of the high potential of DNA metabarcoding to be included in future national marine biomonitoring campaigns.

## 1. Introduction

The spread of non-indigenous species (NIS) poses a significant threat to the diversity and integrity of coastal ecosystems, often leading to profound ecological alterations (Rilov & Crooks, 2009). These species, transported by human activities – whether voluntary or involuntary - can drastically reshape ecosystems by changing biotic interactions (competition, predation, introduction of parasites and diseases) and/or modifying environmental conditions (Bax et al., 2003; Crooks, 2002; Grosholz, 2002). Invertebrates are the most prevalent European marine NIS (Zenetos et al., 2022). The primary pathway for their introduction is shipping, both commercial and recreational (Clarke Murray et al., 2014; Sardain et al., 2019). With the continued growth of global trade and recreational boating, marinas and ports have become critical hotspots for the introduction of marine and brackish NIS (Bulleri & Chapman, 2010; Connell, 2000). These species hitch rides by encrusting ship surfaces or being carried in ballast water, sometimes colonizing the artificial structures of marinas and ports upon arrival (Carlton, 1985; Drake & Lodge, 2007). Recreational boats can further spread these species that might establish viable populations and become invasive, disrupting local ecosystems, but also incurring significant economic costs (Bailey et al., 2020; Clarke Murray et al., 2012; Hewitt & Campbell, 2010; Sempere-Valverde et al., 2024). The ecological impacts include the displacement of native species, alterations in habitat structure, alterations in the food-webs, and changes in ecosystem functions (Tsirintanis et al., 2022). Economically, these invasions can lead to increased maintenance costs for marine infrastructures, reduced fisheries yield, and the need for costly management actions, including eradication, control and containment efforts (Lovell et al., 2006). Addressing this challenge requires coordinated international efforts to monitor and manage the pathways of introduction, particularly focusing on shipping practices (Miralles et al., 2021).

Strategies to manage NIS are costly and complex. Therefore, implementing monitoring programs is crucial for early detection, recording and managing the impacts of NIS on native ecosystems (Hulme, 2009; Leung et al., 2002; Occhipinti-Ambrogi & Savini, 2003). In the European Union (EU), the Marine Strategy Framework Directive (MSFD) was set in place for all EU Member States to protect and conserve the marine environment (European Parliament and Council of the European Union, 2008). This directive requires member states to develop strategies to achieve Good Environmental Status (GES) by 2020 and provide detailed assessments of their marine environments and the impacts of human activities. Descriptor D2 specifically focuses on NIS and requires each EU member state to monitor their presence and conduct risk assessments (Tsiamis et al., 2019). Consequently, European guidelines for marine NIS assessments have been published for several countries, namely France, Denmark, and the Netherlands (Gittenberger et al., 2023; Jensen et al., 2023; Massé et al., 2023). In Portugal, a first list of NIS was provided for the Azores (Cardigos et al., 2006) and Madeira (Canning-Clode et al., 2013) archipelagos, and the first overall national assessment was published in 2015 for coastal ecosystems in mainland Portugal, Madeira and the Azores, based on literature review and sampling surveys conducted in coastal ecosystems. A total of 133 NIS were identified, of which 85 (ca. 64%) were marine and brackish invertebrates (Chainho et al., 2015). The Reports of the Working Group on Introductions and Transfers of Marine Organisms (WGITMO) for Portugal updated these numbers annually, with the most recent report indicating the occurrence of 211 NIS in the Portuguese coastal areas, with 53 invertebrates identified in the Azores and Madeira archipelagos and 119 in mainland Portugal (ICES, 2022). These assessments have been reflected in the reports for D2 within the MSFD reports for Madeira, the Azores, Mainland and Continental Platform, with the latest update available for public consultation to be published in 2025, as a result of the 3^rd^ cycle of implementation of the MSFD (ME, SRMP, SRAAC, 2025). Regular NIS assessments in Portugal have also been published as a result of sporadic research projects in the Azores and Madeira (Castro et al., 2022, 2023; Parretti et al., 2020; Png-Gonzalez et al., 2021) and in mainland Portugal (Chebaane et al., 2023; Ribeiro et al., 2023). Although periodic NIS surveys have been conducted in several recreational marinas, using PVC settlement plates (Souto et al., 2018) and scrapping fixed areas in hard substrates (Afonso et al., 2020; Ribeiro et al., 2023), these assessments are not continuous in time and no consistent methodological approaches nor surveillance areas have been used. Also, in most marine NIS surveys in Portugal, these species have been identified exclusively by morphology-based approaches (Afonso et al., 2020; Chainho et al., 2015a; Chebaane et al., 2023; Encarnação et al., 2024; Piló et al., 2021; Rubal et al., 2021). Although these traditional approaches have been instrumental in assessing the status of NIS in Portugal, they are hindered by several significant drawbacks. Identification of marine NIS using solely morpho-taxonomy is a slow and labour-intensive process that requires phylogenetically-diverse taxonomic expertise, which is becoming increasingly scarce (Hopkins & Freckleton, 2002; Kim & Byrne, 2006). This limitation hampers the ability to provide the rapid responses typically demanded by management authorities. Additionally, these approaches are less effective for the early identification of NIS, which requires detecting species present at low abundances or in spatially restricted areas (Walsh et al., 2018). Early detection improves the chances of a successful containment or eradication of NIS, before the species becomes widely established (Ojaveer et al., 2018; Zanden et al., 2010).

To overcome the above-mentioned challenges, DNA metabarcoding has been widely adopted in biomonitoring programs due to its effectiveness in processing many environmental samples simultaneously, allowing for rapid and accurate biodiversity assessments (Duarte et al., 2021). By extracting DNA from environmental samples, amplifying specific short genomic regions and processing them through high-throughput sequencing (HTS), species can be accurately identified at the molecular level (Cristescu, 2014; Rey et al., 2020; Taberlet et al., 2012). It also overcomes common difficulties experienced during traditional biomonitoring approaches (such as identification of larvae/smaller organisms, specimens at low density or cryptic taxa) while being comparable and complementary to the traditional morpho-taxonomic approaches (Holman et al., 2019; Keck et al., 2022; Pochon et al., 2013). This technique is very valuable in NIS assessments, as it allows for large-scale surveys with a greater sampling effort and, more importantly, it allows the early detection of NIS before their establishment causes significant, lasting impacts (Steinke et al., 2022; Zaiko et al., 2015). In Portugal, only a few studies have used DNA metabarcoding for surveying marine/brackish NIS (Azevedo et al., 2020; Lavrador et al., 2024); and there is yet to be a national assessment of marine invertebrate NIS using this approach.

In this study, we used environmental DNA metabarcoding to address this gap. Specifically, we aim to: 1) perform a comprehensive geographic assessment of marine and brackish invertebrate communities in ten recreational marinas across mainland Portugal, and the archipelagos of Madeira and the Azores; 2) identify potential hotspots of NIS within the studied area; and 3) compare the NIS detected in this study with the most recent NIS lists available (MSFD lists), as well as with morphological and molecular NIS records from the same marinas, to evaluate the efficacy and constraints of DNA metabarcoding for NIS detection. We anticipate that the diversity of marine invertebrate communities will vary significantly between the mainland Portuguese marinas and those in Madeira and the Azores, since they are located in different biogeographical regions. Furthermore, we expect non-indigenous invertebrate species to be more prevalent in mainland marinas due to higher levels of human activity, and a greater number of recreational marinas, which theoretically provide more available niches for NIS establishment. In addition, mainland Portugal is more connected to other coastal regions with NIS established populations, increasing the potential for species to spread through natural or human-assisted means. Due to the higher sensitivity of high-throughput sequencing, we also expect to detect new NIS, that have not been previously recorded in Portugal.

## 2. Materials & Methods

### 2.1 Sampling sites

Sampling was conducted during June and July of 2020 in ten recreational marinas across mainland Portugal, Madeira and the Azores (Fig. 1, Table S1), including Viana do Castelo and Porto Atlântico marinas in the North of Portugal; Costa Nova and Jardim Oudinot marinas in Aveiro; Alcântara and Oeiras marinas in Lisbon; Funchal and Quinta do Lorde marinas in Madeira Island; and Vila Franca do Campo and Ponta Delgada marinas in São Miguel Island, in the Azores archipelago. The distance between marinas from the same regions varies from 3 km in Aveiro to approximately 57 km in the North of Portugal.

**Figure 1.**
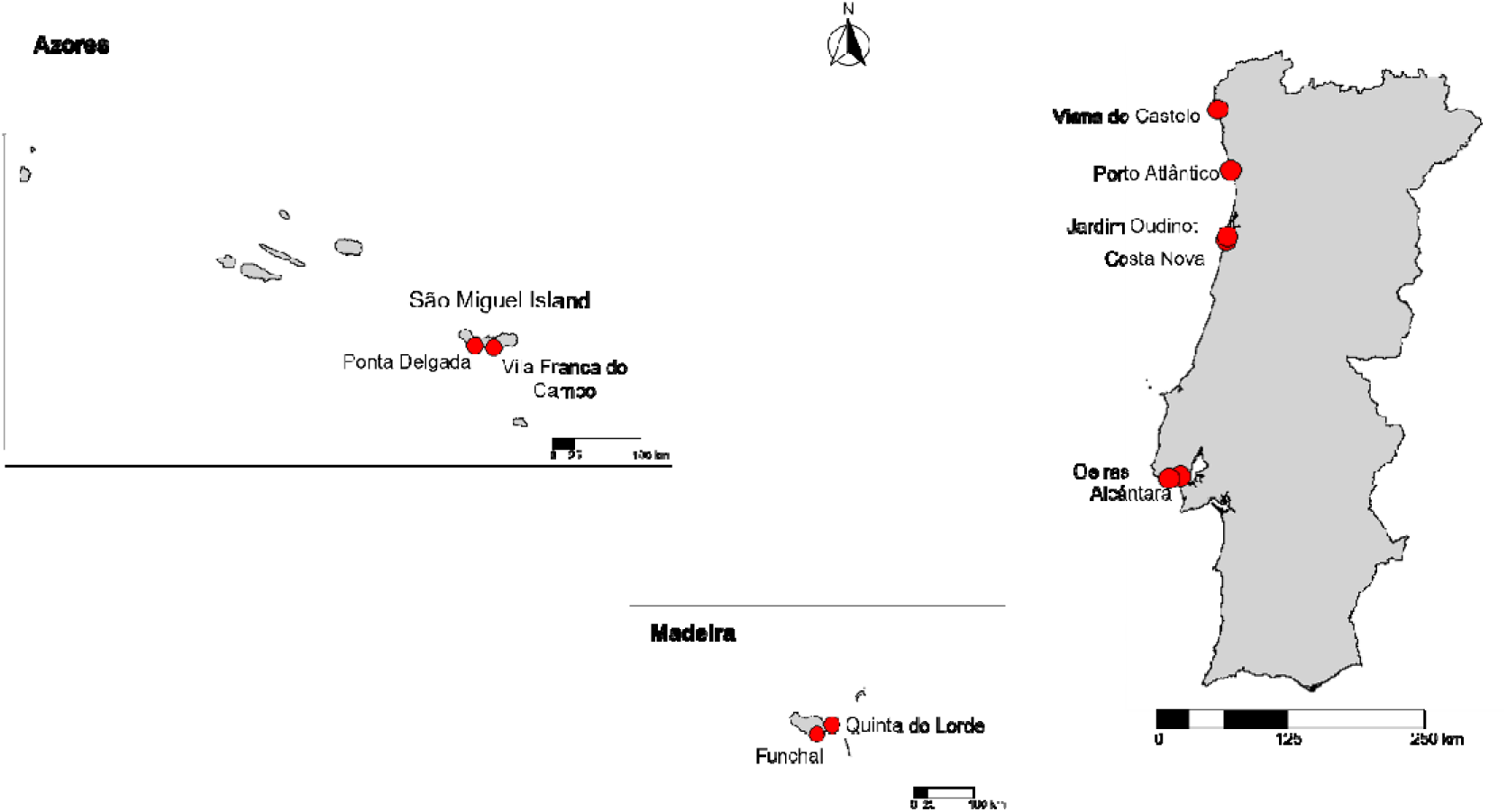
Mainland Portugal and Madeira and Azores archipelagos with red dots displaying the location of the ten sampled recreational marinas.

Jardim Oudinot is the smallest marina, with less than 100 docking stations for boats up to 12 meters in length, while Ponta Delgada is the largest marina with up to 640 docking stations for ships up to 90 meters in length. Overall, the remaining marinas have 100-300 docking stations for ships up to 30 meters.

These ten marinas also differ in distance to the nearest harbour as well as in their position on the coast. In the north of Portugal, the Viana do Castelo marina is located on the northern margin of the Lima River, at approximately 2 km from the Viana do Castelo Port (this port received 195 ships in 2024); while Porto Atlântico marina is located near the mouth of the Leça River, at less than 1 km from the Leixões harbour (which is the second harbour with the highest shipping traffic in Portugal, receiving in 2024 a total of 2.343 ships). In Aveiro, the two marinas differ in their position within the Ria de Aveiro: Costa Nova is more sheltered, while Jardim Oudinot is closer to the sea. Jardim Oudinot is also located closer to the Port of Aveiro (less than 1 km, which received 958 ships in 2024). In Lisbon, the Alcântara marina is located in the Tagus estuary closer to the Port of Lisbon (ca. 1 km), which is one of the busiest ports in Portugal (received 1.132 ships in 2024), while the Oeiras marina is closer to the sea, located at a distance of ca. 15 km of the that port. In Madeira, both marinas have similar dimensions and are near commercial ports (Ramalhosa et al., 2021), with Funchal marina being closer to the Port of Funchal (which received 1.244 ships in 2024). In the Azores, Ponta Delgada marina is larger and within the Ponta Delgada Port (which received 835 ships in 2024), while Vila Franca do Campo marina has 80% less docking capacity and is further away from the commercial port, albeit near a fishing harbour.

### 2.2 Sample collection, processing and DNA extraction

Three types of samples were collected in each marina: 1) fouling communities in hard substrates (e.g. pontoons, ropes and buoys) were scraped into a tray of 22 x 22 cm, and then transferred into zip-lock bags with water from the site; 2) a water sample (1L) was collected at ca. 1-meter depth into polypropylene flasks; and 3) zooplankton was collected using a plankton net (55 µm mesh size, 100 cm length and 40 cm diameter mouth; with oblique hauls of approximately 45 seconds and 3.5-4 meters deep along the water column) into 1L polypropylene flasks. All propylene flasks had been previously washed with 10% bleach and rinsed with MilliQ water. Negative controls of 1L flasks with MilliQ water were used throughout all the workflow. All samples were stored at 4° C for up to 24 h until processing. One sample per substrate type was collected at four different pontoons per marina to ensure spatial representativeness. Zooplankton was not collected in Madeira due to technical constraints.

Organisms collected from hard substrates, along with the water from the zip-lock bags, were sieved through a 500 µm mesh previously bleached (10% bleach). The retained organisms were stored in absolute ethanol at -20 °C. DNA extraction from these organisms was performed using a silica-column based method. They were incubated overnight in a lysis buffer (100 mM NaCl, 50 mM Tris-HCl pH 8.0, 10 mM EDTA pH 8.0, and 0.5% SDS), followed by centrifugation, binding, and purification through silica columns (Lavrador et al., 2024; Steinke et al., 2022). Water for eDNA analysis, water from the negative controls, and zooplankton samples were filtered through 0.45 µm pore size filters (S-Pak Filters, Millipore); DNA was then extracted from half of the filters using the DNeasy PowerSoil Kit (Qiagen), following the manufacturer’s instructions. Negative controls were processed with the DNA extraction procedure to check for contamination of the solutions and labware materials used. These negative controls were also used as templates in subsequent PCR amplification reactions.

### 2.3 Amplicon libraries, high-throughput sequencing (HTS) and bioinformatic pipelines

To create amplicon libraries, the 5’ region of the mitochondrial cytochrome c oxidase I (COI) gene [mICOIintF/LoboR1 primer pair (Leray et al., 2013; Lobo et al., 2013)] and the V4 region of the 18S rRNA gene [TAReuk454FWD1/TAReukREV3 (Lejzerowicz et al., 2015; Stoeck et al., 2010)] were targeted. Sequencing was performed in an Illumina MiSeq® platform at Genoinseq (Biocant, Portugal), using V3 chemistry for paired-end sequencing. Raw reads were quality filtered by minimum length (100 bp for COI and 150 bp for 18S) and trimming of bases (average quality below Q25) (as detailed in Lavrador et al., 2024). Merging of the filtered reads and primer removal was performed using the mothur software (version 1.39.5) (Schloss et al., 2009). mBrave – Multiplex Barcode Research and Visualization Environment (www.mbrave.net; Ratnasingham, 2019) was used to process COI reads, while the SILVAngs database (https://ngs.arb-silva.de/silvangs/; Quast et al., 2013) was used for the 18S reads (also detailed in Lavrador et al., 2024). For COI, reads were uploaded with the following parameters defined - minimum length: 150 bp, maximum length: 313 bp, and reads with QV < 10 were discarded. Following this filtration step, reads were de-replicated, and chimeras were removed. Using a distance threshold of 3%, the remaining reads were clustered into Operational Taxonomic Units (OTUs) that were then matched to the Barcode of Life Database (BOLD) (Ratnasingham & Hebert, 2007) and to specific public datasets of interest using a 97% similarity threshold for taxonomic assignment (details on the used datasets in Lavrador et al., 2024). BOLD uses a Barcode Index Number (BIN) system to group DNA barcode sequences into OTUs that usually correspond to species (Ratnasingham & Hebert, 2013). As certain BINs have more than one species attributed to it, COI records were further curated following this protocol: first, all species were listed per sample and inputted into the BAGS software (Fontes et al., 2021); https://github.com/tadeu95/BAGS) which classifies species records into five grades (A– E) based on the congruency between species assignments and BINs, as well as the quantity of sequences available in the BOLD database. Then, resulting libraries containing species name, respective BIN and attributed grade were exported and analysed. Species and corresponding BIN attributed grades A and B (concordant morphospecies) were maintained; those attributed grade D (insufficient records) were automatically discarded; and those attributed grade C (morphospecies assigned to multiple BINs) and grade E (seemingly discordant morphospecies) were further audited. In short, these records were maintained in case of synonyms or misidentifications that could be solved, while BINs attributed to 3 or more morphospecies and discordances that could not be solved were automatically discarded (details on discordance resolution in Lavrador et al., 2023).

For 18S sequences, in the SILVAngs, the SILVA Incremental Aligner (SINA v1.2.10 for ARB SVN (revision 21008)) (Pruesse et al., 2012) was used along with the SILVA SSU rRNA SEED database for quality control and taxonomic assignment (Quast et al., 2013). Reads with <150 aligned nucleotides, more than 2% homopolymers or 1% ambiguities, or identified as artifacts or putative contaminants were excluded. The remaining reads were dereplicated and clustered into OTUs using VSEARCH (version 2.14.2; https://github.com/torognes/vsearch) (Rognes et al., 2016). BLASTn (2.2.30+; http://blast.ncbi.nlm.nih.gov/Blast.cgi) (Camacho et al., 2009) was used for taxonomic assignment against the non-redundant version of the SILVA SSU Ref dataset (release 138.1; http://www.arb-silva.de). Reads with weak or no classifications, with a “(% sequence identity + % alignment coverage)/2” value below 70, were assigned to the category “No Taxonomic Match”. For subsequent analysis, only taxonomic identifications with >99% similarity threshold were retained. NIS detected exclusively with this marker were further analysed by performing a separate BLASTn with standard settings, against standard NCBI databases, to verify the accuracy of the taxonomic assignments.

For both markers, only reads assigned to species level that belonged to metazoan invertebrate groups were used. Species environment and taxonomic classification were verified using the World Register of Marine Species (WoRMS) database (www.marinespecies.org, accessed from February to April 2024) (WoRMS Editorial Board, 2024). Only marine and brackish invertebrate species were retained.

For NIS identification, final species lists were matched to the most recent Portuguese invertebrate NIS lists from mainland Portugal, Madeira and the Azores (MSFD lists published in 2019 and 2025) (ME, SRMP, SRAAC, 2025; MM, SRMCT, SRAAC, 2019), as well as to an updated European NIS list (Table S2) for the presence of new NIS (details on obtaining the updated European NIS list described in Lavrador et al., 2023).

### 2.4. Data analyses

Venn diagrams were generated to determine the overlap between species and NIS detected in the five different regions (North of mainland Portugal, Aveiro, Lisbon, Madeira and the Azores - https://bioinformatics.psb.ugent.be/webtools/Venn/). Bar graphs and pie charts were used to display the distribution of species and NIS among phyla, for each sampled marina, using GraphPad Prism v8 (GraphPad Software, Inc.). A Principal coordinate analysis (PCoA), using the Jaccard index, was performed to visualize the similarity in community structure among the different marinas and sample types. A permutational multivariate analysis of variance (PERMANOVA) was performed, using 999 permutations, to test the effect of the factors “marina” and “sample type” on the invertebrate community structure. Pairwise comparisons were then performed using the pairwise Adonis package to identify significant differences between individual levels of the factors “sample type” within each marina and “marina” within each sample type. All analyses, including PCoA, PERMANOVA, and pairwise comparisons, were carried out in R using the vegan and pairwiseAdonis packages (Martinez-Arbizu, 2020; Oksanen et al., 2022). A linear regression analysis was performed to assess the relationship between species similarity among samples and geographic linear distance. The geographic distance between marinas was calculated using the distHaversine function in the R package geosphere (Hijmans et al., 2017) (Rstudio v4.0.3). The species similarity matrix was obtained using the Jaccard index calculated in Primer v6.1.16 (Clarke & Gorley, 2006), based on a presence/absence matrix. The final figure illustrating the relationship between geographic distance and species similarity was constructed in R using the package ggplot2 (Wickham, 2016). To evaluate differences in native and NIS richness among regions and between mainland and the islands, one-way ANOVAs were performed in R using the aov() function.

For the comparison between NIS detected in this study and those reported in the MSFD reports, the marinas and locations where these NIS were found, as well as where they had been previously reported, were thoroughly analysed to determine if there were any new NIS records. These analyses were conducted separately for mainland Portugal, Madeira and the Azores, as not all NIS detected in this study are considered NIS in all territories (e.g., *Mytilus* sp., *Clavelina lepadiformis* (Müller, 1776)), and the MSFD NIS lists were also divided by these territories. The MSFD further divided the mainland territory into three regions: A) from Caminha to Peniche (which includes the North and Aveiro marinas of this study); B) Peniche to Lagos (includes the Lisbon marinas); and C) Lagos to Monte Gordo (referred to as “South” in Table S11).

## 3. Results

### 3.1 HTS results

Initial reads for all marinas were 4,917,902 for COI and 4,164,943 for 18S from 222 datasets in total: ((7 marinas x 3 sample types x 4 sampling points) + (1 marina x 2 sample types x 4 sampling points) + (1 marina x 1 sample type x 3 sampling points) + (2 marinas x 2 sample types x 4 sampling points)) x 2 molecular markers. After read merge and primer trimming in the mothur software, 3,350,112 (68%) and 3,276,179 (79%) reads were uploaded into mBrave and SILVAngs, for COI and 18S data processing, respectively. Of these, 1,490,180 reads were taxonomically assigned for COI and 2,627,643 for 18S datasets. Finally, 955,109 and 745,167 reads were assigned to marine and brackish invertebrates (for COI and 18S, respectively), of which 44% and 14% were assigned to NIS (Table S3 and S4). The only negative control that produced sequences was the water eDNA negative control from Ponta Delgada marina, and this was observed exclusively with the 18S marker. This negative control sample was processed using mothur and SILVAngs, following the same protocol as for the other 18S samples. All hits identified in this negative control were subsequently removed from the corresponding water samples. For the remaining analyses, data from both markers were combined.

### 3.2 Differences in species and NIS detected among regions

In total, 645 marine/brackish invertebrates were detected in the ten marinas, of which 40 were NIS (Fig. 2; Table S5 and S6). Regarding each region, the highest number of species was recorded in the north of mainland Portugal (295), followed by the Azores (275), Aveiro and Lisbon (259 each), with Madeira recording the lowest numbers (163). Regarding NIS, the Azores registered the highest number (25), whereas Madeira recorded the lowest (11).

**Figure 2.**
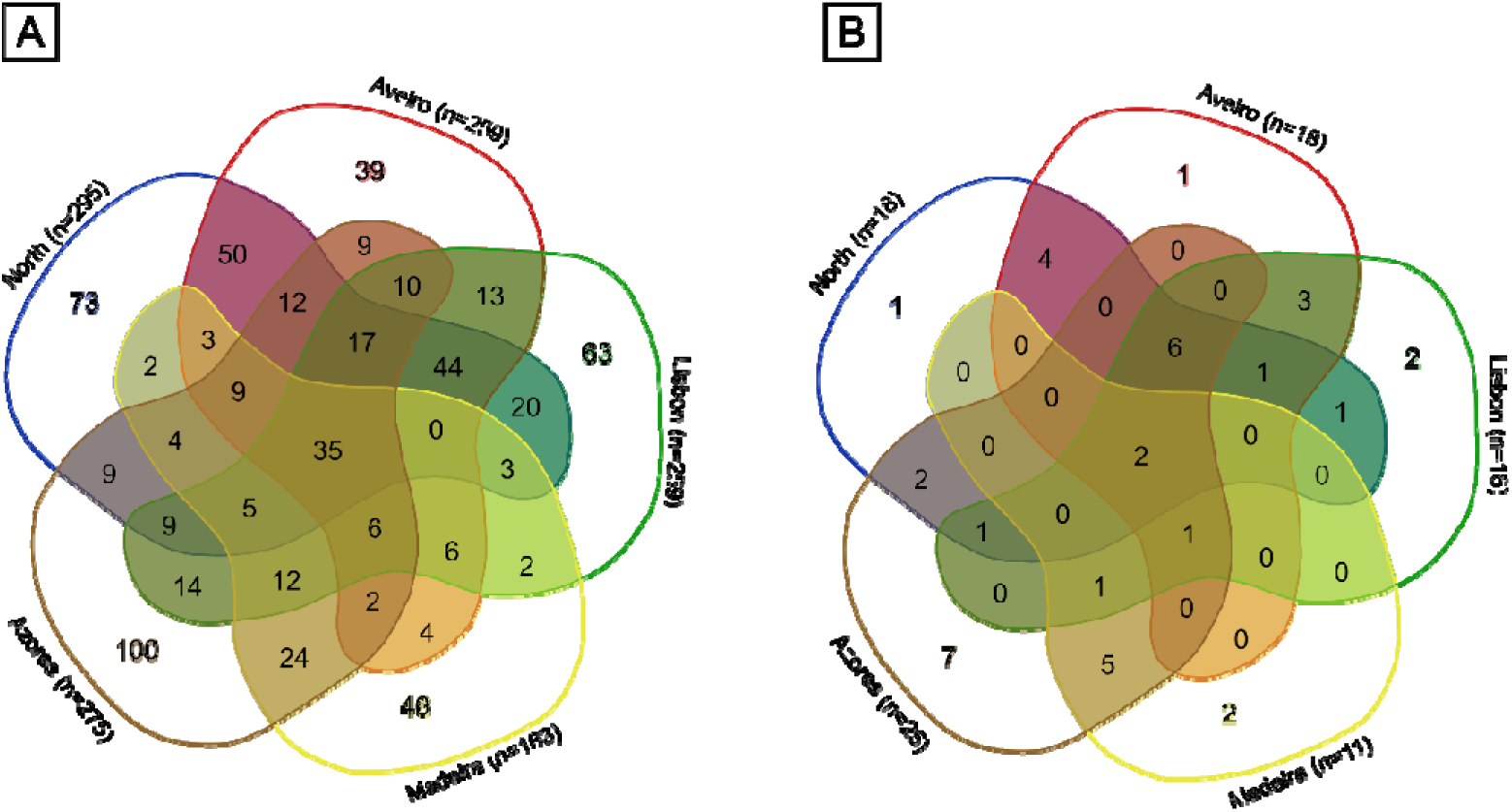
Venn diagrams showing the total (**A**) and non-indigenous species (**B**) detected in the five regions combining data from both marinas of each region.

Analysing the differences between all ten marinas, the highest number of species was recorded in Viana do Castelo (220) (North) and the lowest in Funchal (110) (Madeira). The highest number of NIS was detected in Ponta Delgada (23) (Azores) while the lowest was found at Quinta do Lorde (7) (Madeira). Overall, only 5% of total species and 5% of NIS were detected across all marinas. The highest number of exclusive species and NIS was detected in the Azores, while the lowest were detected in Aveiro and the North of mainland Portugal (in the case of NIS) (Fig. 2). Statistical comparisons showed that the number of native species did not differ significantly between mainland Portugal and Madeira (p=0.208), or between mainland Portugal and the Azores (p=0.875). Similarly, NIS richness did not vary significantly among the regions (p=0.167) or between mainland Portugal and the islands (p=0.275).

Regarding taxonomic composition, the dominant phyla were Annelida, Arthropoda, and Mollusca, with notable regional variations: Annelida was dominant both in Viana do Castelo (25% of the species detected in that marina), and in Madeira (up to 25%); Arthropoda predominated in Aveiro (up to 24%); and Mollusca was the dominant phylum in Lisbon and the Azores (up to 29% in each region), although less detected in Madeira (only up to 10% of species). There were also other distinct patterns such as Rotifera only being detected in the North and Aveiro marinas, and Phoronida not being detected in Madeira nor in the Azores. Ascidiacea was more prevalent in Lisbon, Madeira and the Azores, and Arthropoda was less dominant in the Azores compared to the other four regions (Fig. 3).

**Figure 3.**
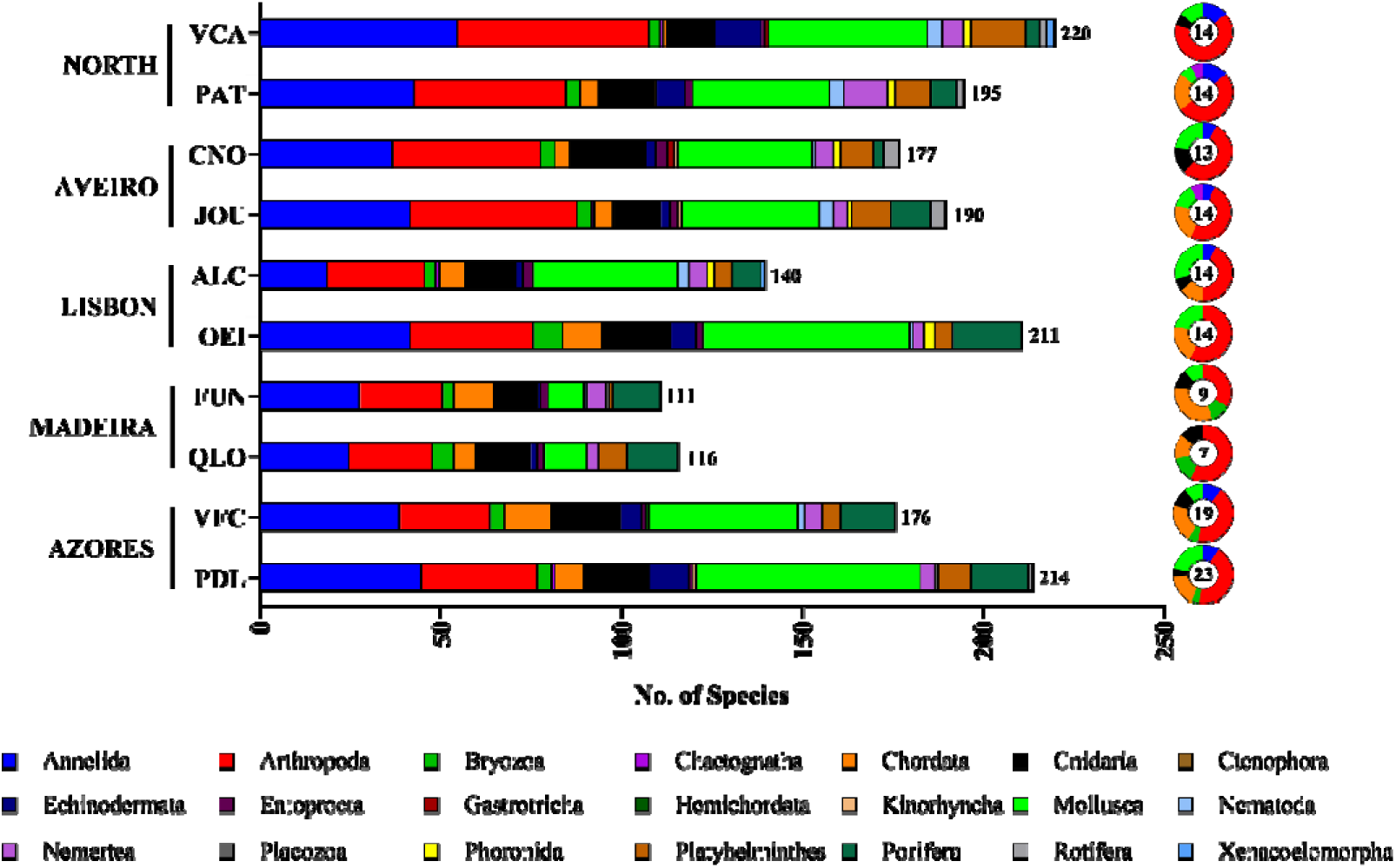
Bar graph displaying the taxonomic composition of the total marine/brackish invertebrate species detected in each marina and pie charts displaying the taxonomic composition of the non-indigenous species detected in each of the marinas with the number of NIS detected in the center of the circle. VCA: Viana do Castelo. PAT: Porto Atlântico. CNO: Costa Nova. JOU: Jardim Oudinot. ALC: Alcântara. OEI: Oeiras. FUN: Funchal. QLO: Quinta do Lorde. VFC: Vila Franca do Campo. PDL: Ponta Delgada.

Most detected NIS belonged to the phylum Arthropoda (33 to 64%). Non-indigenous bryozoans were only detected in Madeira and the Azores; Annelida were mostly detected in the North and in Costa Nova (Aveiro) and no annelid NIS were detected in Oeiras (Lisbon) and the two Madeira marinas. The lowest numbers of non-indigenous Mollusca were detected in both Madeira and Porto Atlântico (North) marinas.

### 3.3 Differences in species and NIS among sample types

The highest percentage of species in total was recovered by zooplankton (63.3%), while the lowest was recovered by water samples (51%). Regarding region, the highest percentage of species were detected in zooplankton samples in the North (69.8%), Lisbon (66.4%) and the Azores (58.9%), while hard substrate samples recovered the highest percentage of species in Aveiro (62.5%) and Madeira (76.1%) (Fig. 4, Table S5). In terms of NIS, zooplankton samples yielded the highest percentage (78.9%), and hard substrate samples the lowest (84.2%). In Aveiro, Madeira and the Azores the highest percentage of NIS was detected in hard substrates (64.7, 90.9 and 83.3%, respectively), while zooplankton samples recovered the highest percentage of NIS in the North and Lisbon (70.6 and 72.2%, respectively) (Fig. 4 and 6, Table S6).

**Figure 4.**
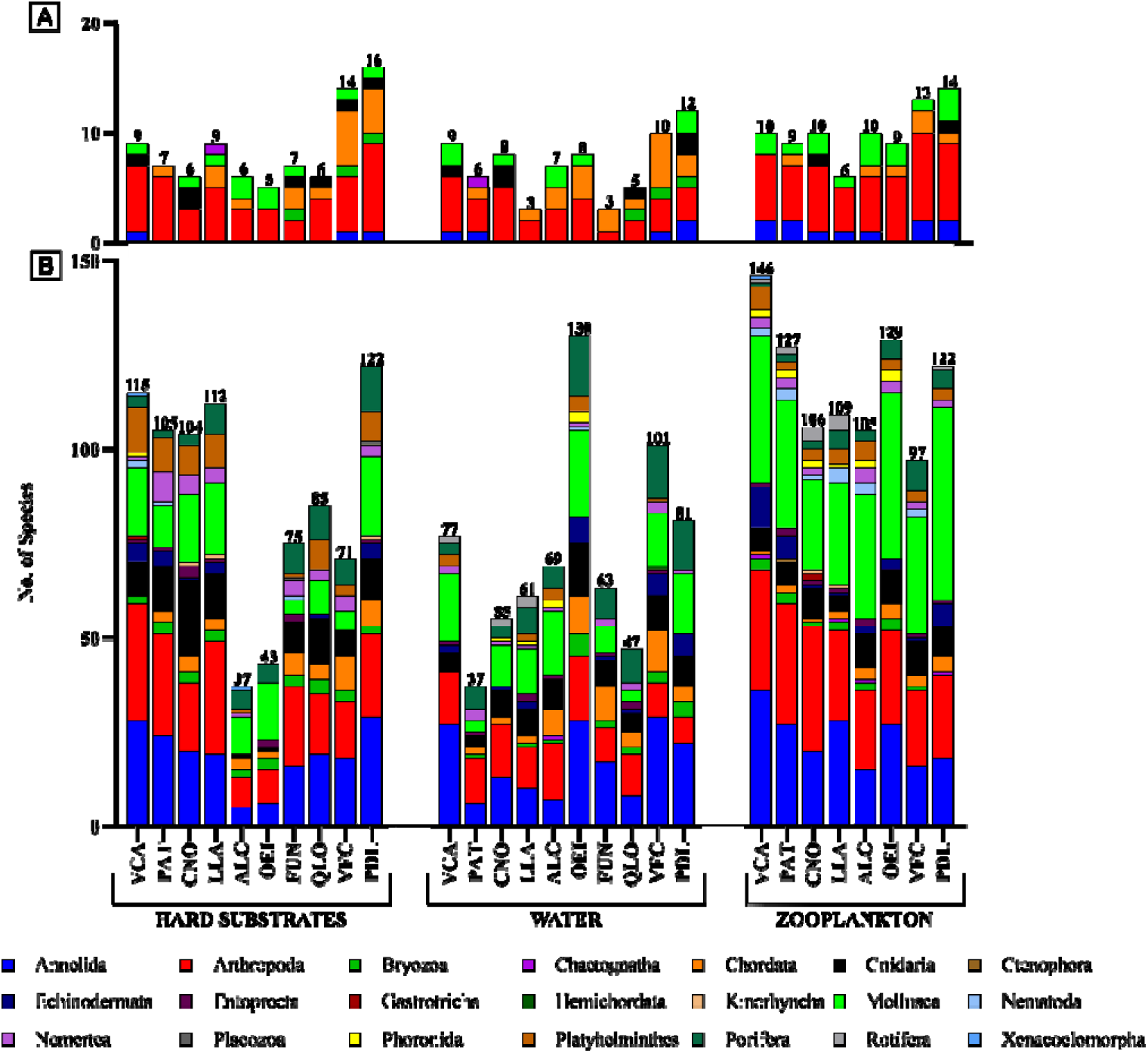
Taxonomic composition of non-indigenous marine/brackish invertebrate species (A) and the total marine/brackish invertebrate species (B) detected by sample type in each recreational marina. VCA: Viana do Castelo. PAT: Porto Atlântico. CNO: Costa Nova. JOU: Jardim Oudinot. ALC: Alcântara. OEI: Oeiras. FUN: Funchal. QLO: Quinta do Lorde. VFC: Vila Franca do Campo. PDL: Ponta Delgada.

Across the marinas, taxonomic composition varied by sample type, but some clear patterns emerged. In hard substrates, Annelida and Arthropoda dominated in most regions, with Mollusca also prominent in the Lisbon marinas. Notably, there were differences between the Aveiro marinas, as Costa Nova was dominated by Annelida and Cnidaria, whereas Jardim Oudinot displayed a more even distribution among Annelida, Arthropoda, and Mollusca (Fig. 4).

In water eDNA, taxonomic composition was more variable across marinas. Annelida, Arthropoda and Mollusca were predominant in Viana do Castelo (North), both Aveiro marinas, and Oeiras (Lisbon). However, in Porto Atlântico (North) and Quinta do Lorde (Madeira), Porifera was one of the most dominant phyla. In the Azores, community composition was more balanced among Annelida, Mollusca and Porifera. Mollusca was the dominant phylum in Alcântara (Lisbon), while Annelida dominated in Funchal (Madeira). Zooplankton samples revealed a general dominance of Mollusca species, coupled with either Annelida or Arthropoda (Fig. 4).

A Principal Coordinate Analysis (PCoA) showed a separation among communities into three groups: 1) North and Aveiro; 2) Lisbon; and 3) Madeira and the Azores. It also revealed a structuring of communities by sample type within these three groups (Fig. 5). The PERMANOVA analysis supported these conclusions, as significant differences were found in the species composition among recreational marinas and sample types (p=0.001, for all factors) (Table S7). Pairwise comparisons revealed that within each marina (in all 10 marinas), community structure significantly differed among the three sample types (p<0.05 for all marinas). On the other hand, for each sample type, comparisons among the ten marinas also displayed significant differences in community composition (p<0.05), with the exception of hard substrates from Quinta do Lorde and Funchal marinas (p=0.051) (Madeira), which were not significantly different, though the p-value was close to the significance threshold (Table S8 and S9).

**Figure 5.**
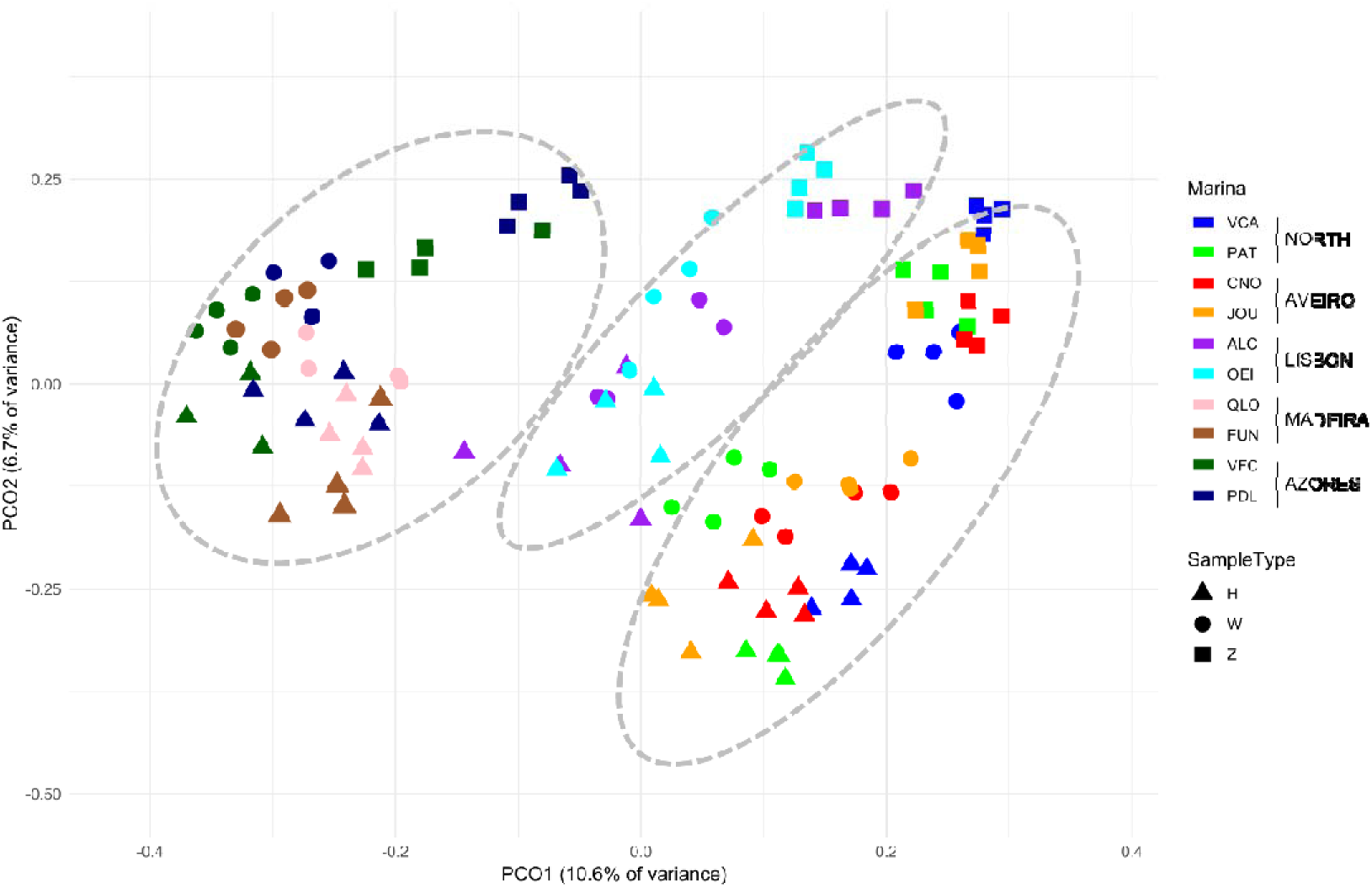
Principal coordinate analysis (PCoA) of sampled communities in the ten recreational marinas, regarding sample type and marina location. Ellipses highlight the separation of communities into three groups: 1) Madeira and Azores marinas (on the left); 2) Lisbon marinas (in the middle); 3) North and Aveiro marinas (on the right). VCA: Viana do Castelo. PAT: Porto Atlântico. CNO: Costa Nova. JOU: Jardim Oudinot. ALC: Alcântara. OEI: Oeiras. FUN: Funchal. QLO: Quinta do Lorde. VFC: Vila Franca do Campo. PDL: Ponta Delgada. H: Hard substrates. W: Water. Z: Zooplankton.

A linear regression analysis also revealed a significant negative correlation between similarity and geographic distance (p<0.001), indicating a decrease in similarity of invertebrate communities as geographic distance increases. The regression model showed that distance explained 42.6% of the variation in the invertebrate community structure (Fig. 6, Table S10).

**Figure 6.**
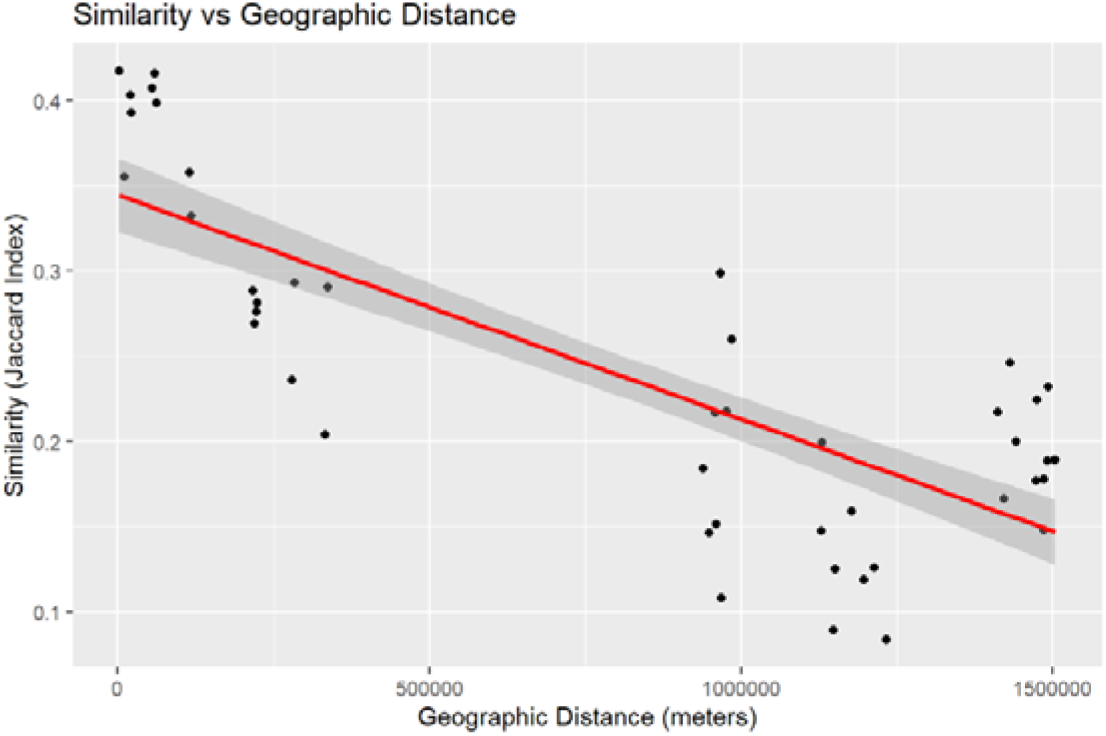
Jaccard index–based similarity among macroinvertebrate species assemblages (including NIS) detected in the three sample types (hard substrates, zooplankton and water eDNA), plotted against straightLline geographic distance between the ten sampled marinas. Red line: fitted linear regression (slope = –1.32 × 10LL kmL¹; R² = 0.62; p < 0.001). Grey shading: 95% confidence interval around the regression line.

### 3.4 Comparison between NIS detected by DNA metabarcoding and those reported in the MSFD lists

In this study, a total of 40 NIS were detected using DNA metabarcoding across mainland Portugal, Madeira and the Azores. Among these, 26 NIS were identified in the six mainland marinas, 11 NIS in Madeira and 25 NIS in the Azores (Fig. 7, Tables S11, S12 and S13).

**Figure 7.**
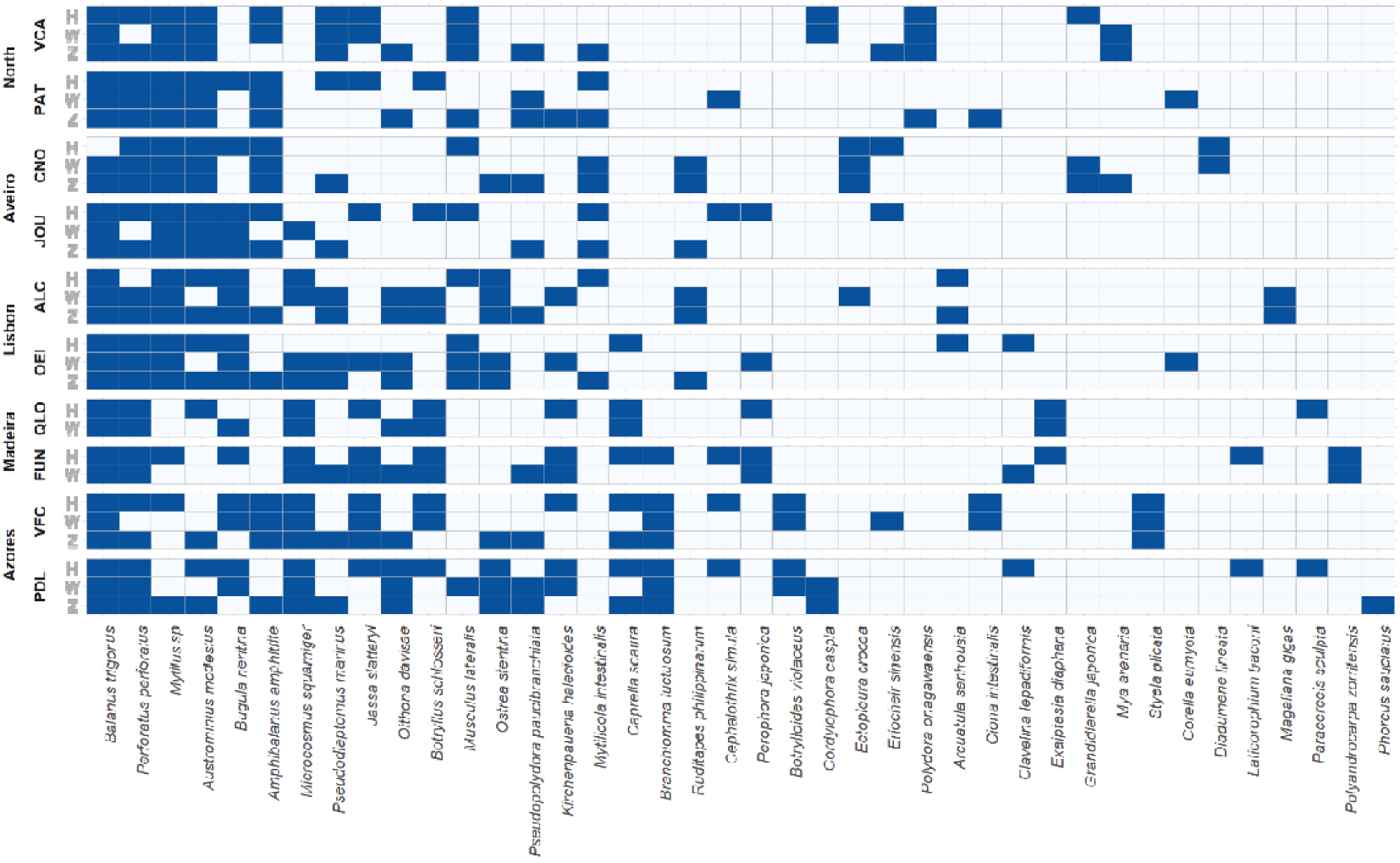
Heatmap of the 40 non-indigenous marine/brackish invertebrate species detected in the ten studied recreational marinas, discriminated by sample. VCA: Viana do Castelo. PAT: Porto Atlântico. CNO: Costa Nova. JOU: Jardim Oudinot. ALC: Alcântara. OEI: Oeiras. FUN: Funchal. QLO: Quinta do Lorde. VFC: Vila Franca do Campo. PDL: Ponta Delgada. Blue colour represents presence of the species in the sample.

When comparing these findings with the most updated national NIS list - the MSFD lists of 2019 and 2025 - new detections and distribution patterns were observed. In mainland Portugal, 9 out of the 25 NIS detected in this study were not classified as NIS in the MSFD reports, but have since been recognized as NIS for the area (ICES, 2022). Examples include the sea anemone *Diadumene lineata* (Verrill, 1869), the tubular hydroid *Ectopleura crocea* (Agassiz, 1862), and the estuarine gammarid *Grandidierella japonica* Stephensen, 1938 (Table S11). Additionally, several NIS showed different distributions than previously reported. For instance, the barnacle *Balanus trigonus* Darwin, 1854, which was previously reported only in Lisbon, has now been detected in all six mainland marinas. Similarly, the Japanese oyster *Magallana gigas* (Thunberg, 1793), formerly reported only in the South of mainland Portugal, has now been found in the Alcântara marina in Lisbon (Table S11).

In Madeira, two NIS – the amphipod *Laticorophium baconi* (Shoemaker, 1934) and *Mytilus* spp. (the entire genus is considered non-indigenous in Madeira) (Segers et al. 2009) - were absent from the MSFD lists but have been since classified as NIS and were detected in Funchal marina (Table S12).

In the Azores, 9 NIS detected in this study were not listed in the MSFD reports for the region. Among them were the oyster *Ostrea stentina* Payraudeau, 1826, the amphipod *L. baconi*, and the mussel *Musculus lateralis* (Say, 1822) (Table S13).

## 4. Discussion

This study provides a comprehensive DNA metabarcoding-based survey of marine and brackish invertebrate communities and NIS in 10 marinas distributed along 3 ecoregions of the lower Lusitanian province, including mainland and island’s locations. Our multi-component approach, integrating three very distinct types of samples (including eDNA), within-site replicate samples, and two target genomic regions, resulted in the detection of 645 marine invertebrate species from 21 phyla, of which 40 were NIS, in a single-time snapshot.

Regarding our first hypothesis, as expected, the diversity and structure of the communities studied varied significantly among marinas. It also varied among different territories within mainland Portugal and between mainland Portugal and islands, as revealed by a PCoA, with the invertebrate communities separating into three main groups: 1) North of mainland Portugal and Aveiro; 2) Lisbon; and 3) Madeira and the Azores. This aligns with previous biogeographic ecoregion and province delimitations, where mainland Portugal is part of a different ecoregion (South European Atlantic Shelf) than Madeira and the Azores (which belong to the Azores, Canaries and Madeira ecoregion), within the Lusitanian Province (Spalding et al., 2007). The separation of the communities from North/Aveiro and Lisbon also corroborates the division of mainland Portugal proposed by the MSFD (ME, SRMP, SRAAC, 2025). The significant negative correlation between community similarity and geographic distance found in this study was also expected, as distant regions often entail different environmental characteristics and ecological interactions, leading to different biological communities (Blanchette et al., 2008; Soininen et al., 2007). Contrary to what would be expected that more non-indigenous invertebrate species would be detected in mainland Portugal, considering that those marinas are more likely to be exposed to higher maritime traffic, all marinas displayed a similar percentage of NIS (6 to 10%). The Ponta Delgada marina in the Azores displayed the highest percentage (10.3%), as well as the highest number of NIS detected (23); and the Quinta do Lorde marina in Madeira displayed the lowest percentage (6%) and number of NIS (7). These lowest numbers of species in Madeira are not unexpected due to the lack of zooplankton samples. The results for the Azores corroborate the invasion patterns identified by Chainho et al. (2015), who found the highest prevalence of NIS in the island habitats and associated it with a higher ecological niche availability due to a lower natural diversity. However, while our study also detected the highest numbers of NIS in the Azores, the overall number of native species was comparable between mainland Portugal and both island regions. Likewise, NIS richness showed similar levels across regions, suggesting that the observed distribution of NIS may instead be influenced by other factors. We also cannot completely rule out the possibility that the Azores receives maritime traffic from a wider range of origins. Finally, the comparison between NIS lists from the MSFD reports and the species detected in our study revealed 9 new NIS records in mainland Portugal, 2 in Madeira and 2 in the Azores (more details in the next sub-section).

### 4.1 NIS records and range expansions in mainland Portugal, Madeira and the Azores

A total of 40 NIS were detected by DNA metabarcoding in mainland Portugal, Madeira and the Azores in the current study. In mainland Portugal, 25 NIS were identified in the 6 sampled marinas, predominantly belonging to Crustacea, Ascidiacea, and Mollusca. Nine of these species were not classified as NIS in the MSFD reports, but were considered NIS based on updated knowledge for these territories (ICES, 2022), such as: the sea anemone *Diadumene lineata*, the tubular hydroid *Ectopleura crocea* and the estuarine gammarid *Grandidierella japonica* (Table S11). In addition, by comparing our findings with the MSFD reports different patterns emerged: 1) *Balanus trigonus* previously reported in mainland only in the Lisbon region, was detected in all 6 mainland marinas; 2) the Japanese oyster *Magallana gigas*, reported only in the South, was now found in the Alcântara marina (Lisbon); 3) *Grandidierella japonica* was found in Viana do Castelo (North) and Costa Nova (Aveiro), whereas it had previously been recorded only in the Algarve (South) (R. Vieira et al., 2024); 4) *Diadumene lineata* was detected in Aveiro, with prior records only in the southern region (Piló et al., 2021; Sousa, 2016); 5) *Ectopleura crocea, Musculus lateralis* and the amphipod *Jassa slatteryi* Conlan, 1990 were newly recorded in Aveiro and Lisbon, while its previous presence was only recorded in the northern region (Azevedo et al., 2020; Lavrador et al., 2024) and in Algarve in the case of *J. slatteryi* (Piló et al., 2021); 6) the copepod *Oithona davisae* Ferrari F.D. & Orsi, 1984 was detected in Lisbon for the first time, with previous records only from northern Portugal (Lavrador et al., 2024); 7) the tunicate *Perophora japonica* Oka, 1927 was detected for the first time in Aveiro and Lisbon, with previous records only for Algarve (Albufeira) in 2019 (Chainho, 2020) and 8) the copepod *Pseudodiaptomus marinus* Sato, 1913, previously recorded only in the North/Aveiro area, was now also detected in Lisbon (Table S11).

These new records suggest the spread of these NIS to new areas; however, further studies are needed to confirm their presence and distribution. Regarding two other NIS detected in our study: the copepod *Mytilicola intestinalis* Steuer, 1902 (a parasite of mussels) had been previously recorded in Portugal (Francisco et al., 2010), but it is unclear if our findings represent new detections, given that the last record was over a decade ago. Considering the report of this species in Galicia (Paul 1983), its presence in Portugal is not entirely surprising and it has in fact been detected in previous metabarcoding studies in Northern Portugal (Hollatz et al. 2017). Being a parasite, it is likely that *M. intestinalis* has been present, probably for several years, but overlooked, which highlights the added value of molecular tools in detecting inconspicuous or underreported species. The polychaete spionid *Polydora onagawaensis* Teramoto, Sato-Okoshi, Abe, Nishitani & Endo, 2013 had its first recorded presence in the North (Lavrador et al., 2024) where it was also detected in this study. Additionally, two species listed as NIS in the MSFD lists and detected in this study are no longer considered NIS: the amphipod *Ampithoe valida* S.I. Smith, 1873 (due to evidence suggesting a ubiquitous distribution (Harper et al., 2022), and the hard clam *Mercenaria mercenaria* (Linnaeus, 1758) (its long history of introductions and the lack of clear historical records have led some authors to propose a possible cryptogenic status (Haydar, 2012).

In Madeira we detected 11 NIS, of which the majority were arthropods and tunicates (Table S12). Of these, a notable case is the first molecular record of the amphipod *Laticorophium baconi* for Madeira (absent from the MSFD lists, but now classified as NIS – Guerra-García et al., 2023). In our study it was detected molecularly in June/July 2020 in Funchal marina, and it has since been identified based on a morphological assessment in Calheta marina (approximately 26 km from Funchal marina) in January 2022 (Guerra-García et al., 2023). Other two interesting cases arose in this region: the tunicate *Botryllus schlosseri* (Pallas, 1766), which was detected in both marinas and listed as NIS in the MSFD reports, has since been classified as cryptogenic (Castro et al., 2022) and, therefore, was not considered a NIS in this study. Conversely, the hydrozoan *Ectopleura crocea*, previously classified as a NIS in Madeira (Chainho et al., 2015), was not detected by DNA metabarcoding in Madeira, despite being found in other marinas included in our study.

In the Azores, 25 NIS were detected, of which 8 were the first molecular records of the hydrozoan *Cordylophora caspia* (Pallas, 1771), the amphipods *Jassa slatteryi*, *Laticorophium baconi*, the mussel *Musculus lateralis*, the copepods *Oithona davisae* and *Pseudodiaptomus marinus*, the oyster *Ostrea stentina* and the polychaete spionid *Pseudopolydora pauchibranchiata* (Okuda, 1937). The barnacle *Austrominius modestus* (Darwin, 1854) is also a first molecular detection for the Azores, although there is some ambiguity surrounding its status as a new record. Historical evidence indicates that this species was already reported in São Miguel in 1903 (Gruvel, 1907). However, its presence has been questioned by several authors (Buckeridge & Newman, 2010; Southward, 1998; Torres et al., 2012). In light of this, we consider this record a new DNA-based detection of *A. modestus* in the Azores. Another interesting case is the hydrozoan *Ectopleura crocea*, a known NIS in the Azores which was detected by DNA metabarcoding in other regions, but not in the Azores. This new data suggests the possible introduction or expansion of these species. However, to verify these findings, extended NIS assessments should be conducted, including detailed morphology-based surveys that would also help to elucidate the population abundance status.

Adding new species to official national NIS databases involves several verification steps, and often identifications obtained through eDNA require validation by morphological assessments due to the limitations of DNA-based identifications (see subsection 4.3). Therefore, some species considered NIS in our study but not present in the MSFD lists might have been known to experts at the time of the reports’ publication; nevertheless, it still needs to be confirmed by morphological assessments. Other discrepancies between MSFD NIS lists and known NIS in the Portuguese territory can be explained by new information on geographical distribution that changes the status of the species (e.g., *Ampithoe valida*); or unknown or recent introductions (e.g., *Musculus lateralis* and *Oithona davisae*). These new records may also represent early detections of NIS in these regions, as DNA metabarcoding can identify species before they are physically observed. For example, the flatworm *Notocomplana koreana* (Kato, 1937) was detected through environmental DNA studies in 2017 at the Port of Vlissingen (Netherlands), but physical specimens were only confirmed three years later (Gittenberger et al., 2023).

Similarly, the invasive snail *Potamopyrgus antipodarum* (J. E. Gray, 1843) was identified through eDNA at a new site in Pennsylvania, USA, in 2018, with physical confirmation following at the same location (Woodell et al., 2021). Comparative studies using both eDNA techniques and morphological assessments have yielded consistent results. For example, *Oithona davisae* was detected in ballast water samples from the Chesapeake Bay (USA) using both DNA analysis and morphological examination (Rey et al., 2019). Accordingly, we recommend conducting additional molecular and morphological assessments to confirm the presence of the newly detected NIS in these regions.

### 4.2 Emerging patterns, range expansions, and discrepancies between different methodologies

Comparing our NIS detections with the most recent morphological and metabarcoding NIS assessments in Portugal revealed several similarities. The barnacles *Amphibalanus amphitrite* (Darwin, 1854) and *Austrominius modestus*, had been found in 2017/2018 in two estuaries close to Viana do Castelo (North) and Costa Nova (Aveiro) marinas (Rubal et al., 2021). In our study, *A. modestus* was also detected in both marinas, while *A. amphitrite* previously found only in the Aveiro estuary, was now also detected in Viana do Castelo (North), which can indicate a range expansion for this species. The tunicates *Ciona intestinalis* (Linnaeus, 1767) and *Botryllus schlosseri* (detected in Porto Atlântico) had also been found and morphologically identified four years earlier in the same area (Leixões harbour) (Azevedo et al., 2020). And of the 27 NIS molecularly detected by Azevedo et al. (2020) (using the COI marker), the barnacle *Austrominius modestus*, the hydrozoan *Bougainvillia muscus* (Allman, 1863), the jellyfish *Clytia hemisphaerica* (Linnaeus, 1767), the sea sponge *Halichondria (Halichondria) panicea* (Pallas, 1766), the barnacle *Perforatus perforatus* (Bruguière, 1789) and the crab *Pilumnus hirtellus* (Linnaeus, 1761) were also detected four years later in our study, suggesting their effective presence. However, we only considered *A. modestus* a NIS, since *Perforatus perforatus* is currently considered a NIS, but only in the Azores. Subsequent to our study, in a 2021/2022 molecular zooplankton assessment in the Lima estuary near Viana do Castelo marina, *Austrominius modestus*, the barnacle *Balanus trigonus*, the copepod *Pseudodiaptomus marinus*, and the polychaete *Pseudopolydora paucibranchiata* were also detected (Moutinho et al., 2023), which suggests that these NIS may have become established in this marina. In Lisbon, the non-indigenous caprellid amphipod *Caprella scaura* Templeton, 1836, initially found outside the Oeiras and Alcântara marinas in 2016 (Afonso et al., 2020), was now detected inside the Oeiras marina four years later. If confirmed through additional morphological and molecular assessments, this could indicate that these species are spreading from the outer areas to the interiors of these marinas or vice-versa. Further ecological studies should be conducted to assess the patterns of spread to and from recreational marinas, as well as to determine potential factors that may lead to these expansions (e.g., recreational boating, natural dispersal).

In Madeira, *Balanus trigonus*, initially found outside Quinta do Lorde marina in 2017, was now detected inside the marina, along with the sea anemone *Exaiptasia diaphana* (Rapp, 1829), which was again detected inside the marina (Diem et al., 2023; Ramalhosa et al., 2019). Both species, along with the bryozoan *Bugula neritina* (Linnaeus, 1758) had also been identified in a 2018 to 2020 assessment in Quinta do Lorde marina (Castro et al., 2023) and were subsequently found in a posterior study in 2021/2022 (Sempere-Valverde et al., 2023), reinforcing the presence of these NIS in Madeira. In the Azores, seven NIS detected in our study - *Botryllus schlosseri*, *Branchiomma luctuosum*, *Bugula neritina*, *Clavelina lepadiformis*, *Kirchenpaueria halecioides*, *Microcosmus squamiger* and *Perforatus perforatus* and *Mytillus* spp.*–* had also been previously identified in Ponta Delgada marina (Castro et al., 2023; Chainho et al., 2015b), reinforcing the efficiency of DNA metabarcoding in the detection of locally established NIS.

However, several species previously identified through morphology-based approaches in these marinas were not detected in our study using DNA metabarcoding. For example, the crab *Cronius ruber* (Lamarck, 1818) found in a 2017/2018 underwater assessment near Funchal and Quinta do Lorde marinas (Schäfer et al., 2019) that was not detected in our study. This discrepancy might be attributed to differences in sampling depth between studies - our analysis focused on surface-level water eDNA and bulk organisms at approximately 1 meter, whereas the other study was conducted at depths ranging from 4 to 11 meters - or to the unsuccessful establishment of this NIS in these marinas. Additionally, the limited availability of zooplankton samples in Madeira could have hindered the detection of larval DNA. The tunicate *Didemnum vexillum* Kott, 2002 was found near Porto Atlântico in 2016/2017, but was not detected by DNA metabarcoding in our study (Azevedo et al., 2020). This species had only been found in two out of twelve months assessed (February and March), which could explain its absence in our samples collected in June of 2020. The barnacle *Amphibalanus improvisus* (Darwin, 1854) and the bryozoan *Tricellaria inopinata* d’Hondt & Occhipinti Ambrogi, 1985 had not been detected in our study but were detected in a subsequent zooplankton assessment in the Lima estuary, close to the Viana do Castelo marina (Moutinho et al., 2023). We believe it would be interesting to perform regular assessments in this marina to determine if these detections represent new introductions or settlements. However, these discrepancies between molecular studies can also be attributed to methodological differences. In fact, in a study we conducted in 2020/2021 at the Viana do Castelo and Porto Atlântico marinas, using five sample types over three seasons (including hard substrates, water and zooplankton, with the second time-point coinciding with the date of this study), *Tricellaria inopinata* and *Ciona intestinalis* were detected in Viana do Castelo in March and June of 2020 (Lavrador et al., 2024), but were not detected in the current study. This discrepancy is likely due to the re-analysis of the same molecular data, as the reads were submitted to mBRAVE and Silvangs five months after the initial study. As reference databases (particularly BOLD) are continually updated, it is common for the same set of sequencing data to yield different molecular identifications when uploaded at different times.

Two notable cases emerged when comparing our results to recent NIS assessments in Portugal. *Laticorophium baconi*, whose most northern record in Portugal is from Sines (2019) (Guerra-García et al., 2023), was detected in this study only in Madeira and the Azores; and *Branchiomma luctuosum*, found in Faro (south of mainland Portugal) in a 2019 assessment (Fernández-Romero et al., 2021), was detected in our study only in Madeira (but only one read in Funchal), and in the Azores where it is considered NIS. As previously mentioned, these differences between studies do not necessarily imply that newly detected NIS in our study have been introduced or have become established in the marinas during the interval between studies. Similarly, the absence of NIS in our study that were previously found in the marinas does not imply their disappearance. Discrepancies in species detection using DNA metabarcoding can be attributed to methodological differences between studies, such as variations in sampling periods (e.g., three sampling periods in Moutinho et al. (2023) *versus* one sampling period in our study, with no overlap), sample types, bioinformatic pipelines, curation methods, and the continuous update of molecular databases (e.g., the reanalysis of molecular data in Viana do Castelo and Porto Atlântico). These factors significantly influence DNA-based species detection across different studies. Therefore, these discrepancies should be carefully analysed, as they do not necessarily indicate the presence or absence of the species detected. With rigorous and standardized monitoring, this data can help establish first records and track changes in community composition over time.

### 4.3 Strengths and pitfalls of using DNA metabarcoding for NIS assessments

DNA metabarcoding offers significant advantages for NIS assessments in marine and coastal ecosystems compared to traditional morphological methods. It enables quick and accurate identifications of a wide range of species from environmental samples, provides higher taxonomic resolution, and is particularly effective for the early detection of NIS. It also enables the detection of small species that are often challenging to detect through visual assessments, such as the copepod parasite *Mytilicola intestinalis*, which was also detected in this study. This allows for a faster and timely implementation of management efforts regarding biological invasions (Duarte et al., 2021; Elbrecht et al., 2017; Westfall et al., 2020).

However, this technique also has its limitations. Namely, the incompleteness of reference libraries which can lead to false negatives, particularly for marine invertebrate NIS in Europe (Duarte et al., 2019), where only ca. 42% of species have COI records in BOLD, and 56% of those are discordant (Lavrador et al., 2023; Vieira et al., 2021). Parallelly, in the case of the survey of native species, the high regional heterogeneity in the levels of completeness of reference libraries (Leite et al., 2020) can also be problematic. These issues can be mitigated using multiple molecular markers (such as the use of COI and 18S in this assessment) and through the curation of molecular results or the use of curated databases. This approach has been shown to increase the number of NIS detected in metabarcoding datasets and to increase species-level identifications in other taxonomic groups (Duarte et al., 2021; Gold et al., 2021; Lavrador et al., 2023; Magoga et al., 2022). For instance, initially in this study (after COI data curation) 45 NIS had been detected. However, further curation of 18S-derived NIS records led to the exclusion of five records, reducing the number of NIS detected in this study to 40, including the exclusion of what would have been a new NIS in the Azores. In this study, three different sample types were used, as it has been shown that community composition in environmental DNA metabarcoding studies varies significantly with sampled substrate/sample type, and the use of a single substrate underestimates species diversity and richness including NIS (Koziol et al., 2019; Lavrador et al., 2024). This was also observed in this study, as the pairwise comparisons revealed significantly different community compositions both among sample types and marinas. In addition, species that are solely detected in the water column through eDNA metabarcoding may never be able to establish themselves in the ecosystems, and, therefore, they should not be immediately included in the NIS lists. Morphological assessments, while more time-consuming and expertise-demanding, allow for estimates of abundance and provide visual confirmation of NIS when identification to species-level is possible.

## 5. Conclusions

Our study showcases several important strengths of multi-component DNA metabarcoding surveys: i) high-throughput and accurate identifications across a broad set of phyla (providing reference libraries are properly curated) ii) parallel multi-community surveys (benthos and zooplankton) using distinct and unique (eDNA) sample types. iii) cross-validation of morphology-based detections, particularly important for taxonomically challenging and highly diverse marine invertebrate communities; iv) highly-sensitive molecular detections of new NIS introductions or range expansions; v) co-detection of native taxa and NIS, enabling contrasting the spatial spread of the two types of organisms, and explore insights into native versus NIS interactions, as for example the possible displacement of closely related natives and the greater or lower susceptibility of hosting communities to NIS introductions.

Several weaknesses persist, mainly regarding incompleteness of reference libraries, molecular detection failures, or inability to provide abundance estimates, but these are expected to be gradually mitigated, as the reference libraries are further completed and curated, and the DNA-technology efficiency globally improves.

We recommend that national monitoring campaigns incorporate DNA-based techniques, ideally combining frequent molecular assessments, which are easier, quicker, and effective with periodic morphological and ecological assessments to provide abundance estimates, community status, and confirm molecularly detected NIS. This approach would enable a continuously updated inventory of species and NIS in national recreational marinas, expanding our knowledge of marine/brackish invertebrate communities in Lusitanian coastal ecosystems, facilitating studies on invasion pathways and about the role of biotic and abiotic factors structuring coastal invertebrate communities.

## Declaration of competing interest

The authors declare that they have no known competing financial interests or personal relationships that could have appeared to influence the work reported in this paper.

## Funding

Funding: This work was funded by the project “NIS-DNA: Early detection and monitoring of non-indigenous species (NIS) in coastal ecosystems based on high-throughput sequencing tools” funded by the Portuguese foundation of Science and Technology (FCT, I.P. under the reference PTDC/BIA-BMA/29754/2017); by the “Contrato-Programa” (https://doi.org/10.54499/UIDB/04050/2020), funded by national funds through the FCT I.P. and in the scope of the project UID/50027-Rede de Investigação em Biodiversidade e Biologia Evolutiva; S.D. was granted financial support by the FCT I.P. (https://doi.org/10.54499/CEECIND/00667/2017/CP1458/CT0001); and A.S.L. (https://doi.org/10.54499/UI/BD/150871/2021) was supported by the Collaboration Protocol for Financing the Multiannual Research Grants Plan for Doctoral Students with financial support from FCT I.P. and the European Social Fund under the Northern Regional Operational Program—Norte2020. P.R. was financed by Fundação para a Ciência e Tecnologia (FCT) through the strategic project [UIDB/ 04292] granted to MARE UI&I. I.A. and J.P.M. were financed by FCT through two PhD grants (SFRH/BD/145746/2019 and SFRH/BD/77539/2011, respectively). P.C., I.A. and J.P.M. were also supported by the FCT through the project UID/04292-Centro de Ciências do Mar e do Ambiente, awarded to MARE and through project LA/P/0069/2020 (https://doi.org/10.54499/LA/P/0069/2020) granted to the Associate Laboratory ARNET.

## Acknowledgments

The authors would like to thank Romana Santos for allowing the use of the laboratory she is responsible for at the Faculty of Sciences of the University of Lisbon, and Judite Alves for allowing the use of the Genetics Laboratory at the National Museum of Natural History and Science at the University of Lisbon, thus enabling us to carry out the laboratory work.

## Author Contributions

Ana S. Lavrador: Formal analysis, Data curation, Investigation, Visualization, Writing - original draft, Writing - review & editing. Inês Afonso: Investigation, Writing - review & editing. Paula Chainho: Investigation, Writing - review & editing. Ana Cristina Costa: Investigation, Writing - review & editing. João Paulo Medeiro: Investigation, Writing - review & editing. Manuela Isabel Parente: Investigation, Writing - review & editing. Paola Parretti: Inves-tigation, Writing - review & editing. Patrício Ramalhosa: Investigation, Writing - review & editing. Ronaldo Sousa: Writing - review & editing. Pedro E. Vieira: Investigation, Writing - review & editing. Filipe O. Costa: Conceptual-ization, Funding acquisition, Writing - review & editing. Sofia Duarte: Conceptualization, Formal analysis, Funding acquisition, Investigation, Project administration, Visualization, Writing - original draft; Writing - review & editing.

## Notes

### Competing Interest Statement

The authors have declared no competing interest.

